# Network of Interactions between the Tumor Necrosis Factor Superfamily Members and Small S100 Proteins

**DOI:** 10.64898/2026.01.08.698460

**Authors:** Victoria A. Rastrygina, Evgenia I. Deryusheva, Alexey S. Kazakov, Andrey S. Sokolov, Maria E. Permyakova, Ekaterina A. Litus, Vladimir N. Uversky, Eugene A. Permyakov, Sergei E. Permyakov

## Abstract

Tumor Necrosis Factor Superfamily (TNFSF) comprises 20 members of membrane/soluble signaling proteins regulating cell survival, cell proliferation/differentiation, and innate/adaptive immunity. Targeting signaling of TNFSF members is used clinically to treat several autoimmune and oncological diseases, and bone loss. They and their cognate receptors are in clinical trials as targets for treatment of autoimmune, inflammatory, oncological and other diseases. Recently, some representatives of S100 family of pleiotropic calcium-binding proteins were shown to interact with TNFSF members TNF and TRAIL, thereby suppressing their activity. In this work, we explored selectivity of interactions between soluble forms of 13 TNFSF members and 21 non-fused S100 proteins using surface plasmon resonance spectroscopy. A total of 27 interactions were found between CD70, CD30L, 4-1BBL, TWEAK, APRIL, LIGHT, VEGI and AITRL and Ca^2+^-loaded forms of S100A1/A2/A4/A5/A6/A12/A16/B/P proteins, with equilibrium dissociation constants from 2 nM to 24 μM. Removal of calcium leads to disruption of the interactions. Molecular docking indicates presence of well-conserved binding sites of the both interaction partners. Mutagenesis of S100P evidences involvement of its ‘hinge’ region in binding of CD30L, VEGI and AITRL, as well as F89 residue in VEGI recognition. The revealed network of interactions is potentially important for regulation of the cellular communication mediated by TNFSF/S100 proteins, which could be exploited for targeted therapy of socially significant diseases.

## 1. Introduction

Tumor Necrosis Factor Superfamily (TNFSF) of cytokine-like proteins comprises at least 20 members (including TWE-PRIL [1]; 18 genes) that signal through specific subsets of 29 receptors in humans, thereby affecting fundamental aspects of cell functioning, regulating innate and adaptive immunity [2-4]. The founding member of TNFSF, Tumor Necrosis Factor (TNF) [5], is a potent pro-inflammatory protein that has proven to be one of the most successful targets for treatment of autoimmune diseases (rheumatoid arthritis, Crohn’s disease, psoriasis, *etc.* [2]), with global market for its inhibitors exceeding US$42 billion (The Business Research Company). Monoclonal antibodies to another therapeutically important TNFSF member, RANKL, are clinically used in treatment of bone loss, bone metastases, and multiple myeloma [6]. Similarly, anti-BAFF antibody is approved for treatment of active systemic lupus erythematosus and lupus nephritis [7]. A fusion protein specific to BAFF/APRIL is also registered for treatment of systemic lupus erythematosus [8]. The antibody-drug conjugates specific to receptors of CD30L or BAFF/APRIL (BCMA) are approved for therapy of lymphomas [9] and multiple myeloma [10], respectively. Other therapeutic modalities using receptors of TNFSF proteins include T cell engagers (BCMA) and chimeric antigen receptor-T cell constructs (BCMA and receptor of 4-1BBL) [11]. Recombinant TNF and circularly permuted TRAIL are approved outside of the US for treatment of soft tissue sarcoma and multiple myeloma, respectively [11]. The success of the drugs that interfere with signaling of the TNFSF members stimulates research into therapeutic potential of this protein family and their receptors, many of which are currently in clinical trials as therapeutic targets [11-13].

TNFSF members are type II transmembrane proteins (except for soluble VEGI) with an extracellular C-terminal domain, termed ‘TNF homology domain’ (THD), which has a ‘TNF-like’ fold (SCOP ID 2000041: β-sandwich, 10 strands in 2 sheets, jelly-roll [14]) [2,3]. Their proteolytic cleavage gives rise to soluble forms of TNFSF members. Both membrane-bound and soluble forms of the cytokines are able to bind to their cognate cell surface receptor(s) via the latter’s cysteine-rich domain(s), forming a hexameric complex between 3 ligand molecules and 3 receptor molecules that activates downstream signaling pathways [2]. The reverse signaling is also possible [15]. Soluble forms of the receptors able to modulate activity of their cognate ligands can be produced by proteolysis or alternative splicing [2,3]. Moreover, cytoplasmic tails of the receptors may or may not contain a death domain, resulting in differences in the downstream signaling pathways. In humans, some of the 29 receptors can interact with more than one of the 20 TNFSF members, and some of the latter can recognize more than one receptor, thereby providing at least 40 individual interactions (may be as many as 194) [2,11,16]. Given the broad expression profile of TNFSF members and their receptors (natural killer cells, T, B and mast cells, eosinophils, monocytes, macrophages, dendritic, epithelial and endothelial cells, *etc.* [2]), they establish a vast network of communications between various cell types, which is important for homeostasis, innate and adaptive immunity. Therefore, they are involved in the processes accompanying a wide range of autoimmune, inflammatory, and oncological diseases [11-13].

Considering medical significance of the TNFSF, knowledge of the factors affecting functional activity of its members is a prerequisite for effective use and development of the therapeutic approaches based on modulation of their signaling pathways. Recently, specific members of the S100 protein family have been shown to alter activity of soluble forms of two TNFSF proteins, TNF and TRAIL, against some cancer cell lines via binding of the S100 proteins to TNF/TRAIL [17,18]. S100 proteins are an evolutionary young family of regulatory calcium-binding proteins (except S100A10, which lacks specificity to Ca^2+^) of vertebrates with broad tissue- and cell-specific expression that are widely involved in a variety of vital physiological processes [19-23]. They contain a pair of EF-hand motifs consisting of two α-helices linked by a Ca^2+^-binding loop: an N-terminal S100-specific EF-hand and a C-terminal canonical EF-hand, connected by a flexible ‘hinge’ region [24,25]. Some S100 proteins also contain distinct Zn^2+^/Cu^2+^-binding sites [26,27]. Human S100 family contains 21 small members (78-113 residues) and 4 long members (146-2849 residues) in which the paired EF-hand domain is fused to a predominantly disordered chain [28]. With the exception of monomeric S100G, S100 proteins tend to form non-covalent homo- or heterodimers (or disulfide-linked dimers, in several cases), as well as higher-order multimeric forms [29-31]. Calcium binding to S100 proteins generally leads to exposure of their hydrophobic residues for target recognition [24]. The localization of S100 proteins both inside and outside cells along with wide range of their targets, including various proteins (enzymes, transcription factors, ion channels, receptors and their ligands, *etc.*), lipids, nucleic acids and N-glycans, allows S100 proteins to affect numerous (patho)physiological processes [20,23,32-34]. Since most S100 proteins, including S100A subfamily, are located in the epidermal differentiation complex (human chromosome 1q21), which encodes many genes expressed in epidermal cells and is rearranged in some cancers [35,36], dysregulation of S100 proteins is associated with skin diseases and cancer [37-41]. Specific S100 proteins are also implicated in many cardiovascular, pulmonary, neurodegenerative, metabolic, inflammatory, autoimmune and other diseases [21,22,42-47]. Some of them are used in diagnostics and/or being tested as therapeutic targets [23,44,48-51].

Since structural modelling and mutational analysis (in the case of S100P-sTRAIL interaction) argue in the favor of conservatism of the S100-binding sites of TNF/TRAIL [17,18], other TNFSF members are expected to be specific to some S100 proteins. To verify this hypothesis, in the present work we examined specificity of soluble forms of 13 TNFSF members to 21 non-fused S100 proteins by surface plasmon resonance spectroscopy. The identified network of interactions between representatives of TNFSF and S100 family points out complexity of regulation of their activities, which should be taken into account in the therapeutic interventions aimed at their signaling.

## 2. Materials and methods

### 2.1 Materials

The samples of recombinant soluble forms of the human TNFSF members used in SPR experiments are listed in Table S1 (excluding TNF, TNF-β and TRAIL). With the exception of CD70/APRIL/LIGHT/OX40L and FasL, which were produced in insect and human cells, respectively, they were produced in *E. coli*. Recombinant tag-free human S100A1/A2/A3/A4/A5/A6/A7/A7L2/A8/A9/A10/A11/A12/A13/A14/A15/A16/B/P/G/Z proteins and S100P mutants F89A and Δ42–47 (lacks PGFLQS sequence in the ‘hinge’ region) were produced in *E. coli* as described in [52-57]. Recombinant tag-free human CaM and human PV were produced in *E. coli* and purified according to [58,59]. HSA was from Merck KGaA (Darmstadt, Germany). Protein concentrations were determined spectrophotometrically as described in [60] (see also Table S1 and ref. [54]).

HEPES, sodium hydroxide and hydrochloric acid were purchased from PanReac AppliChem (Darmstadt, Germany). Sodium chloride was from Helicon (Moscow, Russia). Calcium chloride, EDTA and TWEEN 20 were from Sigma–Aldrich, Inc. (St. Louis, MO, USA). Sodium acetate, EDAC, sulfo-NHS and ethanolamine were bought from Bio-Rad Laboratories, Inc. (Hercules, CA, USA). All solutions were prepared using ultrapure water (Simplicity 185 system from Merck KGaA, Darmstadt, Germany).

### 2.2 Affinity of TNFSF members to S100 proteins

The interactions between soluble forms of TNFSF members and S100 proteins at 25°C were studied by SPR spectroscopy using ProteOn™ XPR36 system and ProteOn GLH sensor chip (Bio-Rad Laboratories, Inc.), essentially as previously described for soluble forms of TNF and TRAIL [17,18]. TNFSF members (30-50 μg/mL in 10 mM sodium acetate pH 4.5 buffer) were immobilized on the sensor chip surface (up to 5,000-14,500 RUs) by amine coupling (EDAC/sulfo-NHS), followed by blocking of the remaining activated amine groups with 1 M ethanolamine solution. Analyte (62.5 nM to 16 μM) in a running buffer (10 mM HEPES-NaOH, 150 mM NaCl, 0.05% TWEEN 20, 1 mM CaCl_2_ or 5 mM EDTA, pH 7.4) was passed through the SPR sensor for 300 s (flow rate of 30 μL/min), after which the sensor was washed with the buffer for 200-2,400 s. The ligand was regenerated with 20 mM EDTA pH 8.0 solution for 100-200 s.

The double-referenced SPR sensograms were approximated mostly using a heterogeneous ligand model:

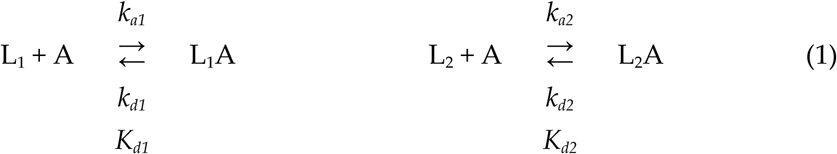

where L_1_ and L_2_ denote two populations of the ligand binding one analyte molecule (A); *k_d_* and *K_d_* correspond to kinetic and equilibrium dissociation constants, respectively. Alternatively, the SPR sensograms were described using either a one-site binding model:

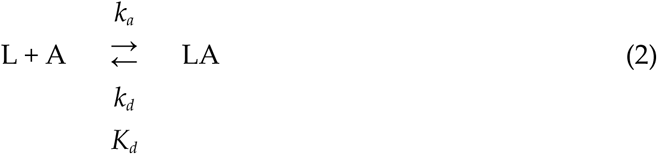

or a two-state model:

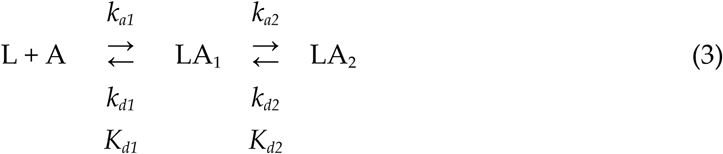

where LA_1_ and LA_2_ correspond to two distinct states of the ligand-analyte complex. The kinetic/equilibrium dissociation constants were calculated using ProteOn Manager™ v.3.1 software (Bio-Rad Laboratories, Inc.) for each analyte concentration and the estimates were averaged (standard deviations are shown, n = 2-9).

### 2.3 Structural modeling of the complexes between TNFSF members and S100 proteins

Structural modelling of the complexes of soluble forms of TNFSF members with S100 proteins was performed within rigid body approximation using ClusPro docking server *[61]*, as described previously [54]. The tertiary structures of the TNFSF members and Ca^2+^-loaded S100 dimers were either taken from PDB [62] (including complexes with their binding partners) or predicted using AlphaFold 3 [63] (https://alphafoldserver.com, accessed 1 June 2025) in the cases where direct structural information was missing or unresolved residues were present in PDB structures (Table S2). For each TNFSF-S100 pair, 10 docking models were built, and the contact residues included in at least 5 models were considered as residues of the probable binding site. The contact residues were mapped onto the amino acid sequences of the binding partners aligned using Clustal Omega [64] (https://www.ebi.ac.uk/jdispatcher/msa/clustalo, accessed 1 June 2025). The predicted tertiary structures of the complexes were drawn with PyMOL v.2.5.0 molecular graphics system [65] (https://pymol.org, accessed 1 June 2025).

## 3. Results and discussion

### 3.1. Selectivty of the interactions between TNFSF members and S100 proteins

Screening of 21 recombinant human small S100 proteins (S100A1/A2/A3/A4/A5/A6/A7/A7L2/A8/A9/A10/ A11/A12/A13/A14/A15/A16/B/P/G/Z) for affinity to 13 recombinant soluble forms of human TNFSF members (Table S1) at a physiological Ca^2+^ level of 1 mM by SPR spectroscopy (TNFSF members are immobilized on the SPR sensor chip via amino groups) revealed 27 specific ligand-analyte interactions, represented as analyte concentration-dependent SPR sensograms with characteristic association-dissociation patterns (Figures 1-3). Since in most cases the dissociation phase is a combination of fast and slow processes, the SPR sensograms were mainly described by the heterogeneous ligand model (1) (Figures 1-3, Table 1), which has been previously successfully used to describe SPR data on S100 binding to soluble TNF/TRAIL [17,18]. In the several cases accompanied by lower SPR signals, the sensograms were approximated with the simplest one-site binding model (2) or the two-state model (3) (Figures 2-3, Table 1). The estimates of equilibrium dissociation constants, *K_d_*, range from 2 nM (TWEAK-S100A12 interaction) to 24 μM (LIGHT-S100A2) (Table 1, Figures 4-5). Similar *K_d_* values were previously reported for soluble TNF (2 nM – 0.7 μM [18]) and TRAIL (0.16-1.8 μM [17]). The range of *K_d_* values for the TNFSF-S100 complexes overlaps with the *K_d_* values reported for complexes of cell-bound members of the TNF receptor superfamily with their ligands (0.01 nM - 19 nM [66]) and for complexes of the S100 proteins with extracellular domains of their receptor, RAGE (2 nM - 23 μM [67]).

**Figure 1.**
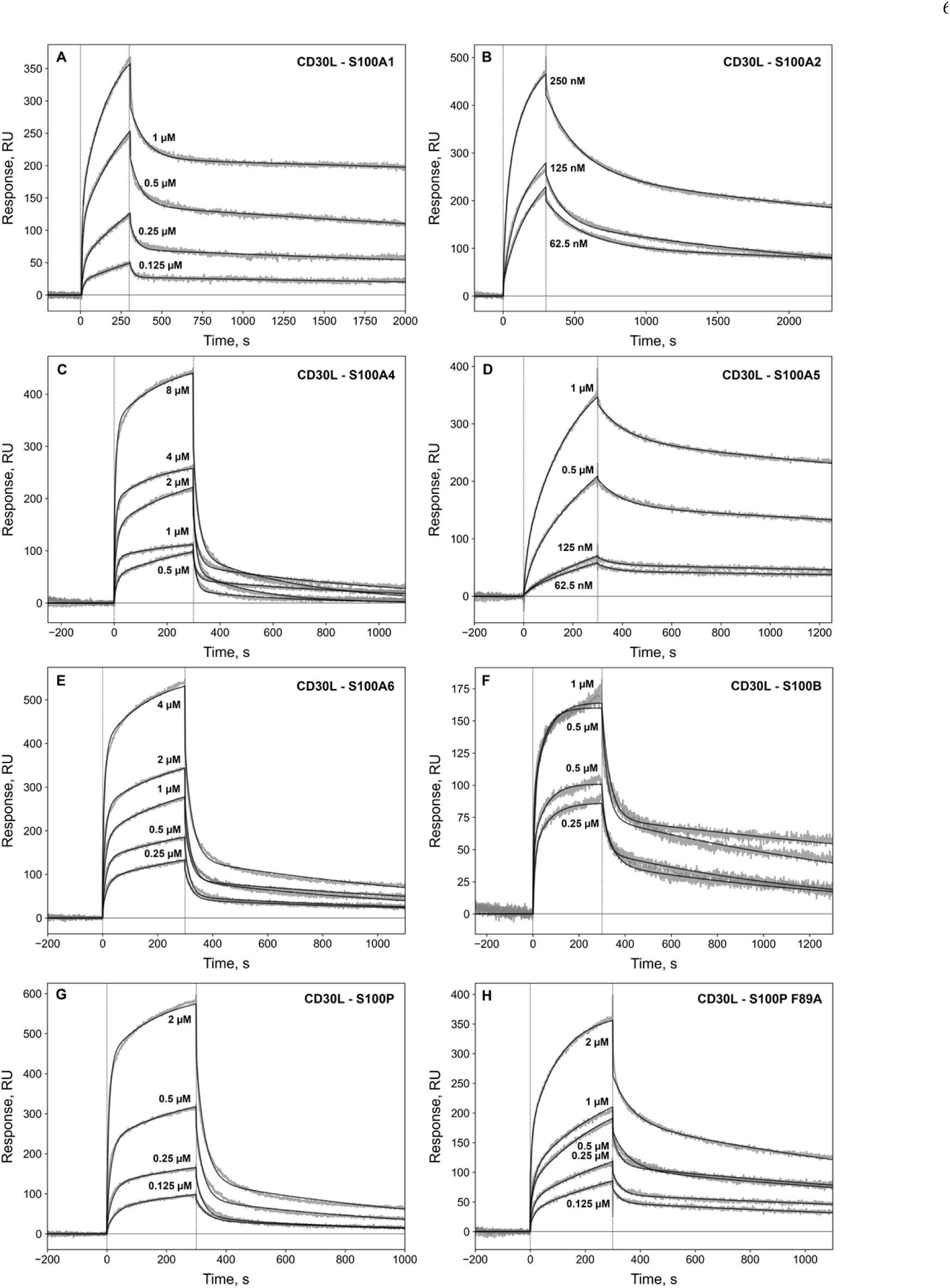
Kinetics of CD30L interaction with Ca^2+^-bound S100A1/A2/A4/A5/A6/B/P proteins or S100P F89A at 25°C, monitored by SPR spectroscopy (*gray*), and theoretical curves (*black*; see Table 1). CD30L is immobilized on the SPR sensor by amine coupling; concentrations of the S100 proteins are indicated for the SPR sensograms. The association phase is indicated by the vertical dashed lines.

**Figure 2.**
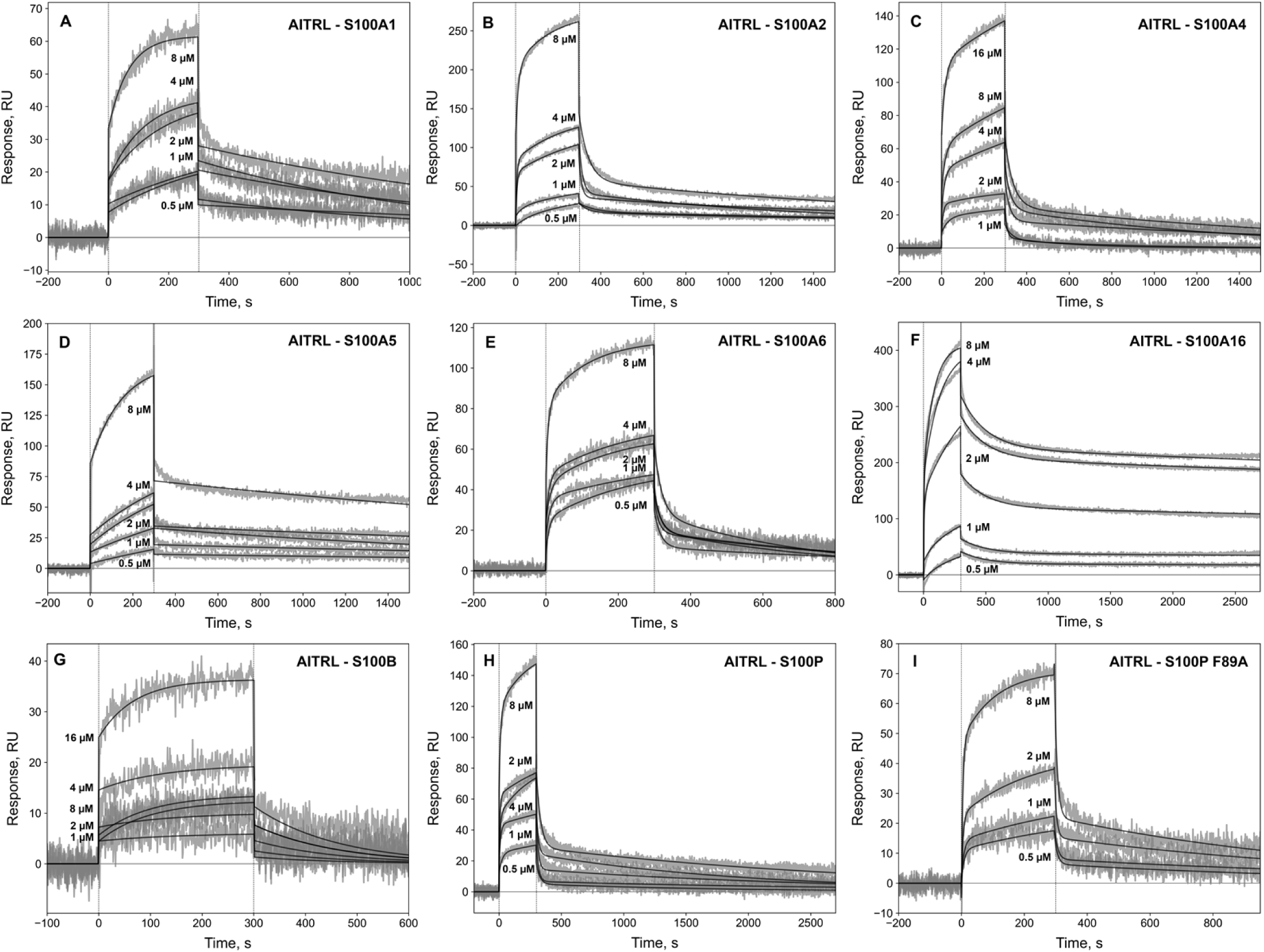
Kinetics of AITRL interaction with Ca^2+^-bound S100A1/A2/A4/A5/A6/A16/B/P proteins or S100P F89A at 25°C, monitored by SPR spectroscopy (*gray*), and theoretical curves (*black*; see Table 1). AITRL is immobilized on the SPR sensor by amine coupling; concentrations of the S100 proteins are indicated for the SPR sensograms. The association phase is indicated by the vertical dashed lines.

**Figure 3.**
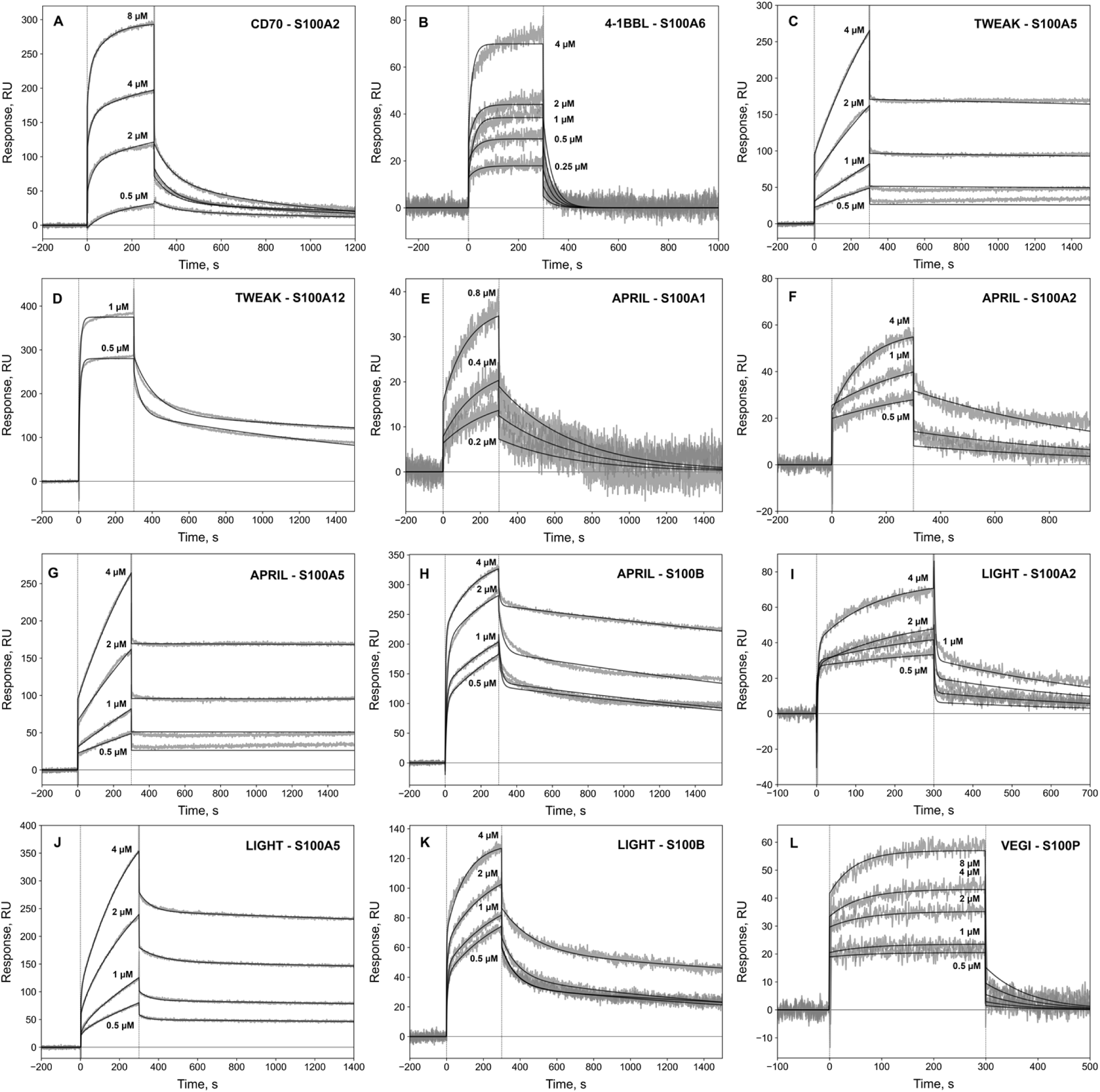
Kinetics of interaction of CD70 (panel **A**), 4-1BBL (**B**), TWEAK (**C**, **D**), APRIL (**E**-**H**), LIGHT (**I**-**K**), VEGI (**L**) with Ca^2+^-bound S100A1/A2/A5/A6/A12/B/P proteins at 25°C, monitored by SPR spectroscopy (*gray*), and theoretical curves (*black*; see Table 1). The TNFSF members are immobilized on the SPR sensor by amine coupling; concentrations of the S100 proteins are indicated for the SPR sensograms. The association phase is indicated by the vertical dashed lines.

**Figure 4.**
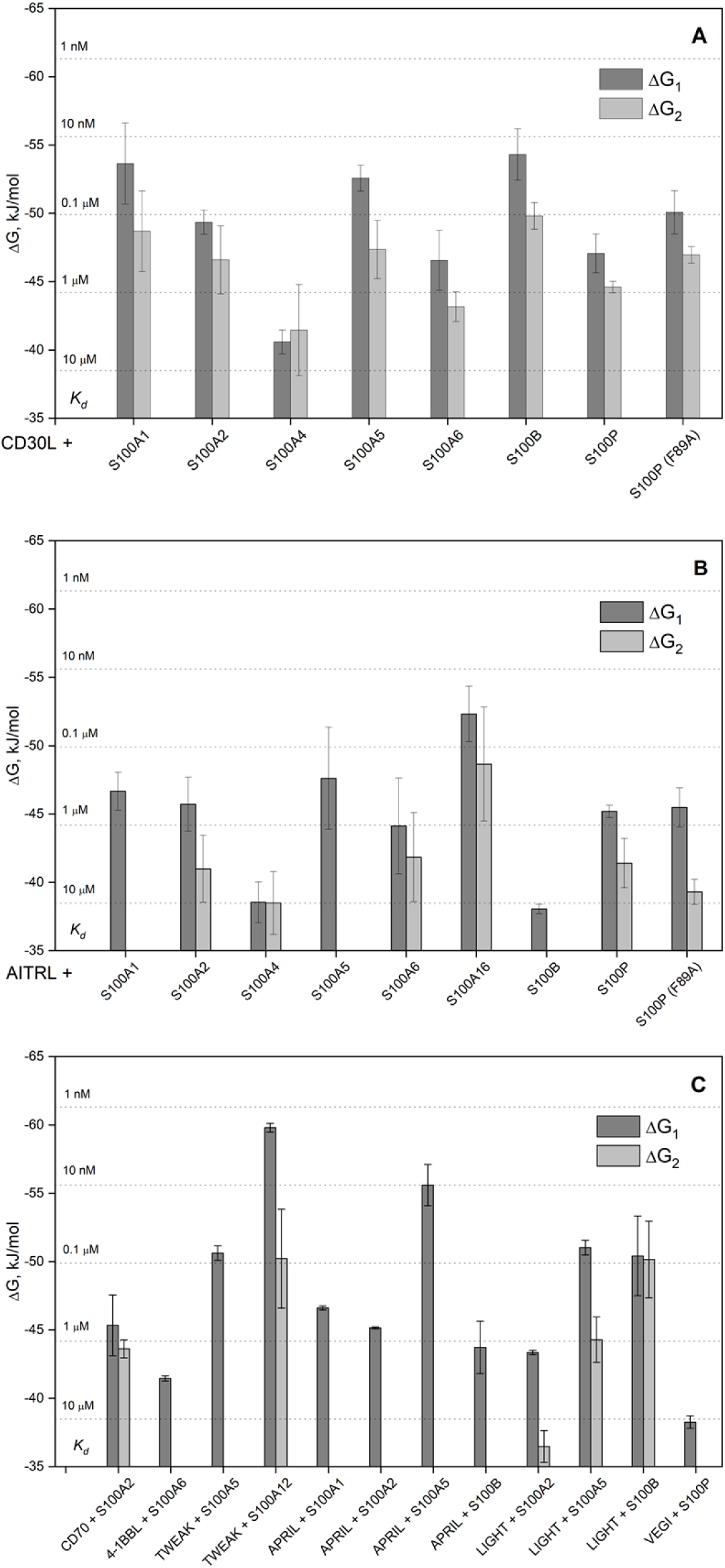
The free energy changes accompanying the binding of Ca^2+^-loaded S100 proteins to TNFSF members at 25°C, calculated from the SPR estimates shown in Table 1: ΔG_i_ = −RT ln(55.3/*K_di_*), i=1,2. The *K_d_* values are indicated by the horizontal dotted lines. See also Figure 5.

**Figure 5.**
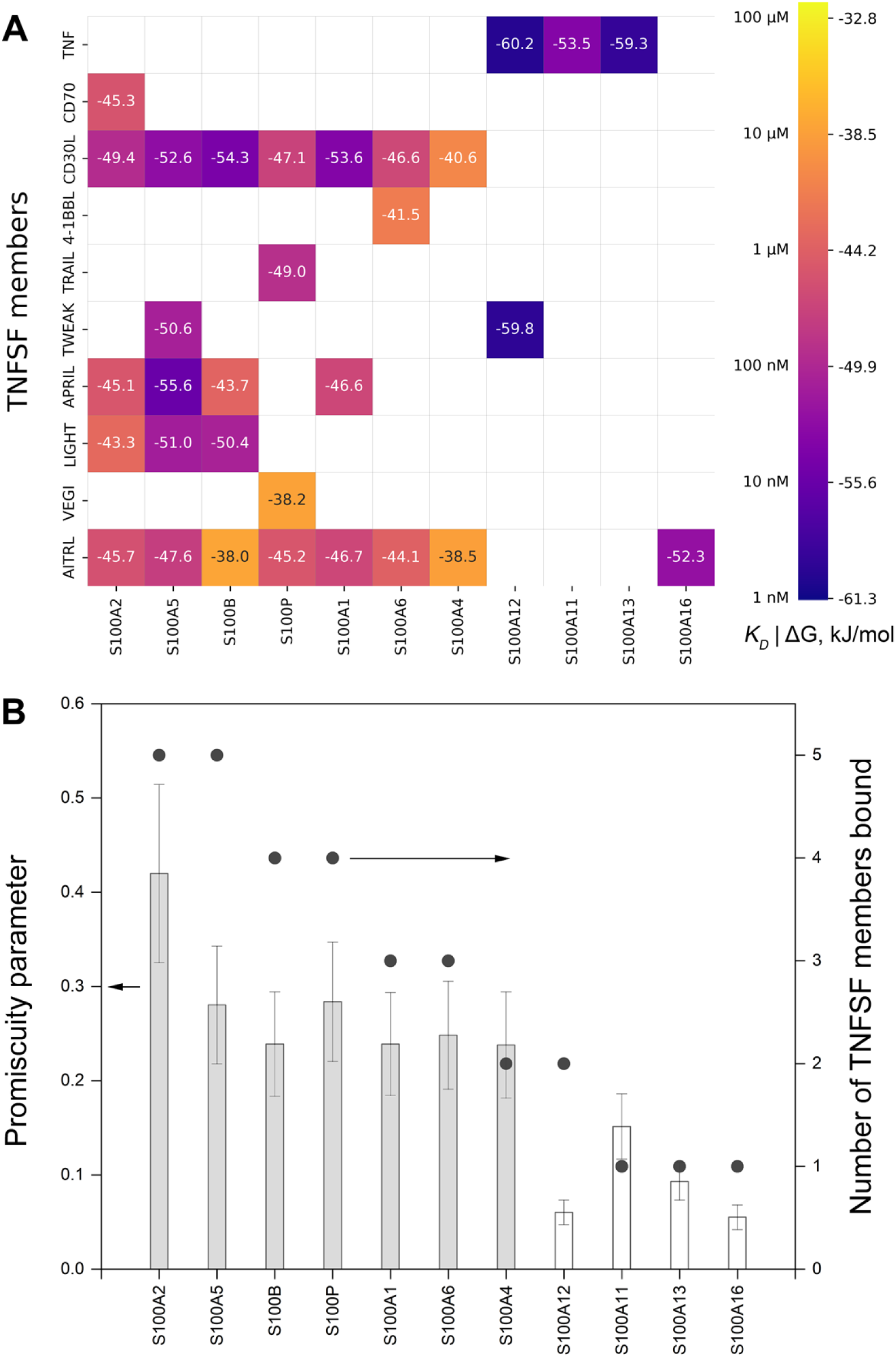
(**A**) Heat map of the highest affinity of soluble forms of TNFSF members for S100 proteins calculated from the SPR spectroscopy data (Table 1, Figure 4) and reported for soluble TNF/TRAIL [17,18]. The lowest ΔG value (kJ/mol) is given for each S100-cytokine interaction. (**B**) Correlation between the promiscuity parameter of S100 proteins [71] (columns) and number of TNFSF members specifically recognized by a S100 protein according to SPR data (filled circles). The columns for “promiscuous” and “orphan” S100 proteins [72] are colored gray and white, respectively.

**Table 1.** The parameters of interaction between soluble forms of TNFSF members (used as a ligand) and Ca^2+^-loaded S100 proteins (analyte) at 25°C, as determined from the SPR data (Figures 1-3), within the heterogeneous ligand model (1), one-site binding model (2) (marked by *) or two-state model (3) (marked by **). See also Figures 4, 5; n/a, not applicable.

| *TNFSF member* | *S100 protein* | $k_{d1}, s^{-1}$ | $K_{d1}, M$ | $k_{d2}, s^{-1}$ | $K_{d2}, M$ |
| --- | --- | --- | --- | --- | --- |
| CD70 | S100A2 | $(7.97 \pm 3.70) \times 10^{-4}$ | $(7.63 \pm 4.85) \times 10^{-7}$ | $(1.24 \pm 0.02) \times 10^{-2}$ | $(1.29 \pm 0.24) \times 10^{-6}$ |
| CD30L | S100A1 | $(1.25 \pm 0.62) \times 10^{-4}$ | $(3.05 \pm 2.10) \times 10^{-8}$ | $(2.47 \pm 1.60) \times 10^{-2}$ | $(2.25 \pm 1.54) \times 10^{-7}$ |
| | S100A2 | $(3.88 \pm 3.52) \times 10^{-4}$ | $(1.29 \pm 0.32) \times 10^{-7}$ | $(1.69 \pm 1.64) \times 10^{-2}$ | $(4.81 \pm 2.94) \times 10^{-7}$ |
| | S100A4 | $(6.71 \pm 1.66) \times 10^{-2}$ | $(4.43 \pm 1.08) \times 10^{-6}$ | $(2.38 \pm 1.31) \times 10^{-3}$ | $(4.51 \pm 3.34) \times 10^{-6}$ |
| | S100A5 | $(1.75 \pm 0.11) \times 10^{-4}$ | $(3.53 \pm 0.94) \times 10^{-8}$ | $(1.31 \pm 0.39) \times 10^{-2}$ | $(3.34 \pm 1.82) \times 10^{-7}$ |
| | S100A6 | $(8.04 \pm 1.03) \times 10^{-4}$ | $(4.64 \pm 2.58) \times 10^{-7}$ | $(4.39 \pm 0.53) \times 10^{-2}$ | $(1.59 \pm 0.48) \times 10^{-6}$ |
| | S100B | $(3.59 \pm 1.09) \times 10^{-2}$ | $(1.08 \pm 0.29) \times 10^{-7}$ | $(6.64 \pm 2.65) \times 10^{-4}$ | $(1.95 \pm 0.96) \times 10^{-8}$ |
| | S100P | $(1.27 \pm 0.23) \times 10^{-3}$ | $(3.40 \pm 1.31) \times 10^{-7}$ | $(3.50 \pm 0.48) \times 10^{-2}$ | $(8.58 \pm 1.03) \times 10^{-7}$ |
| | S100P (F89A) | $(4.60 \pm 0.66) \times 10^{-4}$ | $(1.03 \pm 0.44) \times 10^{-7}$ | $(2.86 \pm 1.18) \times 10^{-2}$ | $(3.33 \pm 0.57) \times 10^{-7}$ |
| 4-1BBL | S100A6* | $(2.73 \pm 0.06) \times 10^{-2}$ | $(3.03 \pm 0.16) \times 10^{-6}$ | n/a | n/a |
| TWEAK | S100A5* | $(3.29 \pm 0.49) \times 10^{-5}$ | $(7.57 \pm 1.14) \times 10^{-8}$ | n/a | n/a |
| | S100A12 | $(3.52 \pm 2.33) \times 10^{-4}$ | $(1.86 \pm 0.17) \times 10^{-9}$ | $(1.71 \pm 1.20) \times 10^{-2}$ | $(1.40 \pm 1.08) \times 10^{-7}$ |
| APRIL | S100A1* | $(2.48 \pm 0.03) \times 10^{-3}$ | $(3.79 \pm 0.16) \times 10^{-7}$ | n/a | n/a |
| | S100A2* | $(1.22 \pm 0.02) \times 10^{-3}$ | $(6.84 \pm 0.15) \times 10^{-7}$ | n/a | n/a |
| | S100A5* | $(4.85 \pm 1.96) \times 10^{-6}$ | $(1.10 \pm 0.45) \times 10^{-8}$ | n/a | n/a |
| | S100B** | $(6.69 \pm 2.78) \times 10^{-2}$ | $(1.40 \pm 0.70) \times 10^{-6}$ | $(2.77 \pm 0.81) \times 10^{-4}$ | n/a |
| LIGHT | S100A2 | $(2.71 \pm 0.13) \times 10^{-1}$ | $(2.38 \pm 0.76) \times 10^{-5}$ | $(1.81 \pm 0.05) \times 10^{-3}$ | $(1.41 \pm 0.06) \times 10^{-6}$ |
| | S100A5 | $(5.79 \pm 0.76) \times 10^{-5}$ | $(6.44 \pm 0.97) \times 10^{-8}$ | $(1.81 \pm 0.67) \times 10^{-2}$ | $(1.07 \pm 0.47) \times 10^{-6}$ |
| | S100B | $(3.19 \pm 1.20) \times 10^{-4}$ | $(1.21 \pm 0.80) \times 10^{-7}$ | $(1.07 \pm 0.34) \times 10^{-2}$ | $(1.11 \pm 0.76) \times 10^{-7}$ |
| VEGI | S100P* | $(1.23 \pm 0.06) \times 10^{-2}$ | $(1.11 \pm 0.14) \times 10^{-5}$ | n/a | n/a |
| AITRL | S100A1* | $(7.12 \pm 3.34) \times 10^{-4}$ | $(3.98 \pm 1.50) \times 10^{-7}$ | n/a | n/a |
| | S100A2 | $(5.76 \pm 1.53) \times 10^{-4}$ | $(6.29 \pm 3.22) \times 10^{-7}$ | $(4.08 \pm 2.80) \times 10^{-2}$ | $(4.60 \pm 2.79) \times 10^{-6}$ |
| | S100A4 | $(1.49 \pm 1.17) \times 10^{-3}$ | $(1.22 \pm 0.71) \times 10^{-5}$ | $(4.95 \pm 3.20) \times 10^{-2}$ | $(1.07 \pm 0.43) \times 10^{-5}$ |
| | S100A5* | $(2.75 \pm 0.86) \times 10^{-4}$ | $(4.09 \pm 3.22) \times 10^{-7}$ | n/a | n/a |
| | S100A6 | $(1.66 \pm 0.52) \times 10^{-3}$ | $(1.59 \pm 1.21) \times 10^{-6}$ | $(6.16 \pm 1.18) \times 10^{-2}$ | $(3.77 \pm 2.75) \times 10^{-6}$ |
| | S100A16 | $(4.15 \pm 0.10) \times 10^{-5}$ | $(4.43 \pm 2.31) \times 10^{-8}$ | $(5.09 \pm 0.05) \times 10^{-3}$ | $(2.98 \pm 2.48) \times 10^{-7}$ |
| | S100B* | $(6.26 \pm 0.34) \times 10^{-3}$ | $(1.20 \pm 0.12) \times 10^{-5}$ | n/a | n/a |
| | S100P | $(4.75 \pm 1.47) \times 10^{-4}$ | $(6.75 \pm 0.87) \times 10^{-7}$ | $(4.98 \pm 1.63) \times 10^{-2}$ | $(3.50 \pm 1.66) \times 10^{-6}$ |
| | S100P (F89A) | $(9.54 \pm 1.36) \times 10^{-4}$ | $(6.47 \pm 2.51) \times 10^{-7}$ | $(1.18 \pm 0.23) \times 10^{-1}$ | $(7.46 \pm 1.92) \times 10^{-6}$ |

Comparison of the *K_d_* values for the complexes between TNFSF and S100 proteins with their maximum serum/plasma concentrations reported in literature and found in the PeptideAtlas project (Figure 6) indicates a potential interaction *in vivo* in the cases of CD30L binding to S100A1/A2/A5/B, TWEAK-S100A12 and APRIL-S100A5. It is noteworthy that comparison of the blood concentrations of the both reagents in these cases (excluding the interactions with S100A5, for which this information is missing) shows that both situations are possible: an excess of S100 protein over a cytokine (TWEAK-S100A12 interaction) and vice versa (CD30L-S100A1) (Figure 6). Therefore, it is expected that the extracellular S100 proteins may affect activity of the TNFSF members, as exemplified by S100-induced suppression of cytotoxicity of soluble TNF/TRAIL toward human cancer cell lines [17,18], and vice versa. Importantly, local extracellular concentrations of the TNFSF/S100 proteins in tumors or inflamed/damaged tissues are expected to exceed those in the blood, thereby favoring their interactions.

**Figure 6.**
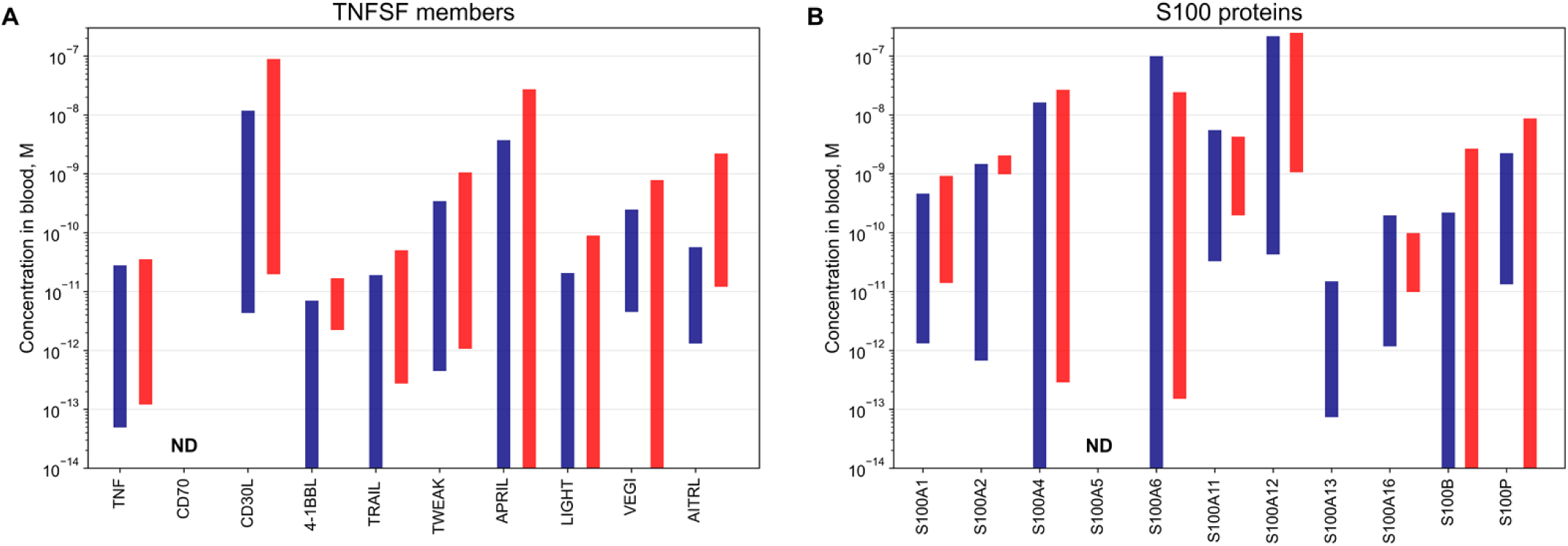
The concentration ranges of the interacting TNFSF (panel **A**) and S100 (**B**) proteins in serum/plasma under normal (blue bars) and pathological (red bars) conditions, according to the immunoassay data found in literature and the PeptideAtlas project (Human Plasma 2023-04 build [75]) (see Supplementary materials SM1). ‘ND’ denotes lack of the data.

The fact that about 10% of the tested cytokine-S100 interactions were positive could be related to notable affinity of EF-hand motifs of S100 proteins for cytokines. This possibility is not supported by the SPR data showing no specific binding of the TNFSF members to recombinant human CaM (Figure S1), a prototypical member of EF-hand superfamily with four EF-hand motifs that recognizes dozens of protein targets [68]. Meanwhile, excess Ca^2+^ is a prerequisite for efficient interaction of the TNFSF members with S100 proteins, since replacement of Ca^2+^ with EDTA [69] leads to disappearance of the SPR effects (Figure S1), which allowed regeneration of the SPR sensor by passage of 20 mM EDTA pH 8.0 solution. The strong calcium dependence of these interactions is in line with previous reports on S100 binding to soluble forms of TNFSF members TNF/TRAIL [17,18] and four-helical cytokines [33,52-54,57,70].

The network of conformation-dependent interactions of TNFSF members with S100 proteins revealed in this work and previously [17,18] is shown in Figure 5A. It includes 31 interactions between ten TNFSF members and eleven s100 proteins with equilibrium dissociation constant values from 2 nM to 24 μM. These interactions are unique due to the considerable evolutionary distance within each group of the binding partners: pairwise sequence identities for the TNFSF members and the S100 proteins are 13%-29% and 25%-61%, respectively. The cross-reactivity of S100 proteins has previously been quantified by introduction of the so-called “promiscuity parameter” [71]. The number of TNFSF members bound per a S100 protein correlates well with the promiscuity parameter (Figure 5B), thus confirming the intrinsic ability of S100 proteins to share protein targets. Notably, some of the S100 proteins that are considered “orphans” [72] (prone to selective target recognition), namely S100A12 and S100A13, exhibit highest affinity for a very limited set of the TNFSF members (TNF/TWEAK; Figure 5), indicating high selectivity and potential physiological significance of these interactions. This observation in the case of TWEAK-S100A12 interaction is consistent with the fact that serum/plasma concentrations of the both reagents (Figure 6) are close to or intersect with the dissociation constant of their complex of 2 nM (Table 1).

Examination of IntAct [73] and BioGRID [74] databases (accessed on May 20, 2025) shows different propensities of the TNFSF members specific to S100 proteins to bind extracellular soluble proteins (the data mainly from large-scale interactome studies): TRAIL and VEGI have no such interaction partners, TWEAK, APRIL, LIGHT and AITRL have 2-4 binding partners, whereas CD70/4-1BBL/CD30L/TNF have 6/11/16/21 interaction partners, respectively. Of these, AITRL and CD30L interact most frequently with S100 proteins (8 and 7, respectively – see Figure 5A), which correlates with a higher propensity to recognize multiple proteins in the case of CD30L. Thus, the ability to CD30L to bind numerous S100 proteins appears to be due to its promiscuous nature, while AITRL is equally prone to recognition of S100 proteins despite its much higher selectivity of protein binding.

Analysis of the binders of the S100-specific TNFSF representatives found in IntAct/BioGRID databases, excluding S100 proteins, shows that only 3 binders contain EF-hands (UniProt IDs Q9H4F8, Q14515 and Q08629). Furthermore, CD30L and LIGHT bind three cytokines, sharing two of them, Bone morphogenetic protein 10 and C-X-C motif chemokine 9, suggesting a possible link between S100 proteins and signaling of these cytokines.

### 3.2. Structural modelling of the complexes between TNFSF members and S100 proteins

Considering low level of amino acid sequence identity within the interacting representatives of TNFSF (13%-29%) and S100 family (25%-61%), their cross-reactivity (Figure 5A) needs a structural explanation. Since calculated pI values of the interaction partners are in the range of 4.5-7.1 (S100) and 6.1-9.8 (TNFSF members, Table S1), their interaction at pH 7.4 cannot be due solely to electrostatic factor. To elucidate structural determinants of this phenomenon, models of the complexes between TNFSF members and Ca^2+^-loaded S100 dimers were constructed in the rigid body approximation using ClusPro docking server *[61]*. The predicted contact residues of S100 proteins (Table S3) are predominantly located at the N- and C-termini, in helices α1 and α3, C-terminal half of helix α4, and in the ‘hinge’ region between helices α2 and α3 (Figure S2A). Notably, these regions of S100 proteins are often involved in recognition of their targets [76], including RAGE [77] and many four-helical cytokines [55,57]. For the TNFSF members, the predicted contact residues (Table S3) are mainly located at the N-terminus, in β-strands A, A’, С, F and H (Figure S2B). Note that some residues of the β-strands A’, С and F are involved in receptor binding [3], suggesting that S100 binding to the TNFSF representatives may affect signaling of the latter’s. Importantly, despite noticeable variation in positions of the contact residues in the aligned sequences of the both interaction partners, locations of their binding sites are well conserved, as illustrated by the models of the least stable (AITRL-S100B) and most stable (TNF-S100A12) complexes (Table 1, Figure 4) presented in Figure 7.

**Figure 7.**
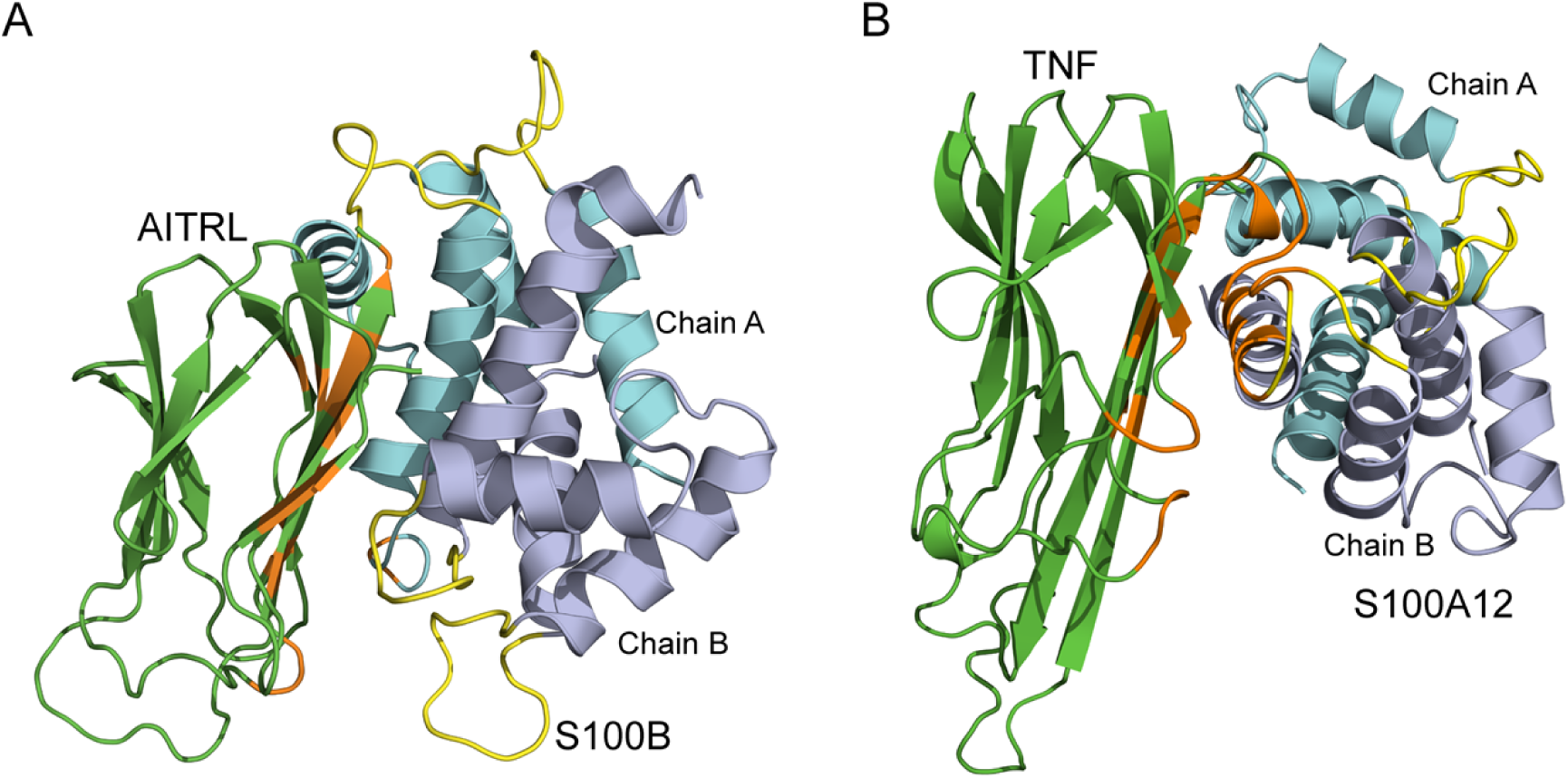
The structural models of the least stable (AITRL-S100B, panel **A**) and most stable (TNF-S100A12, **B**) complexes of TNFSF members (*green*) with Ca^2+^-loaded S100 dimers (*grey* and *cyan*) built using ClusPro docking server *[61]*. The Ca^2+^-binding loops and the contact residues are shown in *yellow* and *orange*, respectively.

To experimentally validate the docking results, S100P mutants Δ42–47 (lacking PGFLQS in the ‘hinge’) and F89A (C-terminal half of helix α4), which were previously shown not to bind TRAIL [17] despite retention of their structural properties [57], were examined for their affinity to CD30L, VEGI and AITRL using SPR spectroscopy. Although residue F89 of S100P protein is predicted to bind all these cytokines (Figure S2A, Table S3), its replacement with alanine caused suppression of S100P affinity to them only for VEGI (Figure S1G), whereas the affinity for CD30L and AITRL remained virtually unchanged (Figures 1-2, 4A, 4B, Table 1). Meanwhile, absence of ‘hinge’ region in S100P Δ42–47 leads to reduced affinity of S100P affinity for CD30L, VEGI, AITRL (Figures S1B, S1G, S1H), although this effect is predicted by molecular docking only for AITRL (Figure S2A, Table S3). Hence, the theoretical contact residues are incompletely correct, which may be due to noticeable structural rearrangements during some S100-TNFSF interactions that are not taken into account in the rigid body approximation used for the predictions. Overall, the experimental data evidence involvement of S100P’s ‘hinge’ in binding to CD30L, VEGI and AITRL, as well as the residue F89 in VEGI recognition, similarly to the previous data on S100P binding to TRAIL [17], many four-helical cytokines [55,57], and RAGE [77].

### 3.3. Intrinsic disorder predspositions of TNFSF members and S100 proteins

To get a potential clue of sequence-encoded features defining differential interactability of TNFSF members and S100 proteins we analyzed the intrinsic disorder predispositions of these proteins. Results of these analyses are summarized in Figures 8 and 9, respectively.

**Figure 8.**
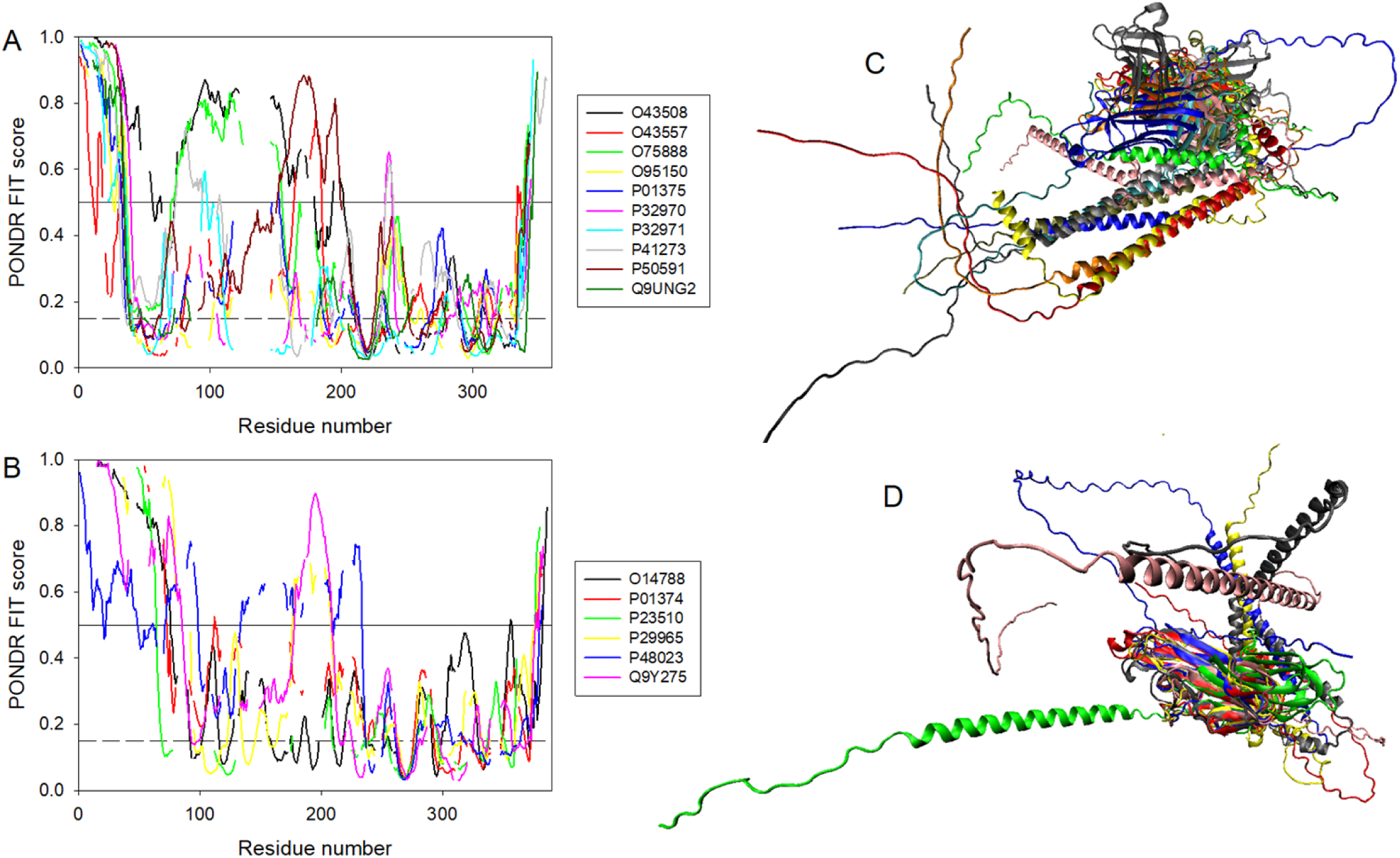
Comparative analysis of structural disorder in the TNFSF members interacting (**A** and **C**) and non-interacting with S100 proteins (**B** and **D**). Aligned intrinsic disorder profiles of the interacting (**A**) and non-interacting cytokines (**B**). Disorder propensity was evaluated by PONDR^®^ FIT [78], and amino acid sequences were aligned using Clustal Omega multiple sequence alignment tool [79] available at https://www.ebi.ac.uk/jdispatcher/msa/clustalo?stype=protein [64]. Aligned 3D structures of the interacting (**C**) and non-interacting cytokines (**D**). 3D structural models for all proteins were generated by AlphaFold [80]. Multiple structural alignment was conducted by SALIGN web server freely accessible to the academic community at http://salilab.org/salign [81].

**Figure 9.**
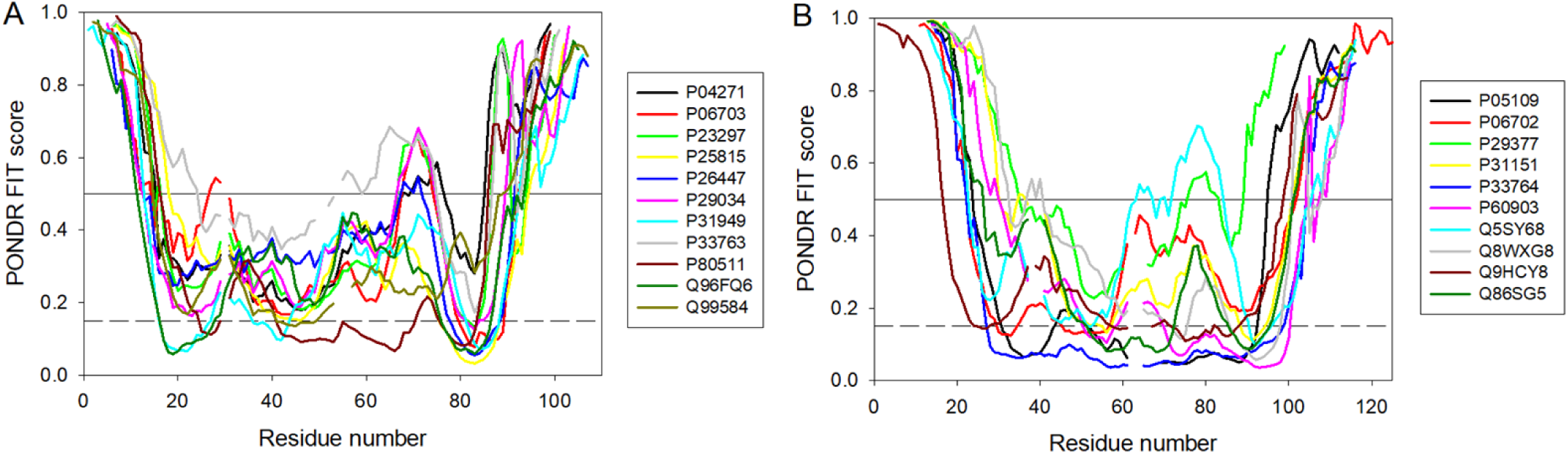
Aligned intrinsic disorder profiles of the S100 proteins capable of interaction with the TNFSF members (**A**) and of the non-interacting S100 proteins (**B**).

Based on the global disorder analysis, the cytokines interacting with S100 proteins were predicted to be a bit less disordered than the non-interacting cytokines (median average disorder scores (ADS): 0.254 vs. 0.299; median predicted percent of intrinsically disordered residues (PPIDR): 16.4% vs. 17.04%). However, the observed differences in the median values between the two groups were not significant, as evidenced by the results of the Mann-Whitney rank sum test (p = 0.704 for median ADS and p = 0.957 for median PPIDR, respectively). Figures 8A and 8B represent aligned disorder profiles of the cytokines. Since amino acid sequences of the TNFSF members are very different, reliability of their sequence alignment is questionable. However, some similarity is observed for the peculiarities of disorder distribution within the C-terminal parts of these proteins (i.e., within the TNF domain region), whereas the N-termini are completely different in this sense. Comparison of Figures 8A and 8B shows that the cytokines interacting with S100 proteins show higher variability in disorder of TNF domains than the non-interacting cytokines. In fact, the aligned intrinsic disorder profiles in the TNF domain containing C-terminal parts in the cytokines interacting with S100 proteins differ from the corresponding regions of the non-interacting cytokines. It appears that in general TNF domains of the S100-interacting cytokines are somewhat more disordered than the C-termini of the non-interacting cytokines (median ADS: 0.189 vs. 0.148; p = 0.092, based on the Mann-Whitney rank sum test) (see also Figures 8A and 8B). Some additional information supporting this notion can be retrieved from analysis of the structurally aligned sets of the cytokines interacting with S100 and the non-interacting cytokines (Figures 8C, 8D). The non-interacting cytokines align along their C-terminal TNF domains reflecting stronger similarities of these domains, whereas the cytokines interacting with S100 proteins show greater variability of their TNF domains, as they align along the long N-terminal helices and show no structural similarity in their C-terminal TNF domains.

Next, we conducted disorder analysis of the S100 proteins interacting and non-interacting with the TNFSF members. Global disorder analysis indicated that the S100 proteins interacting with the cytokines are somewhat less disordered than the non-interacting cytokines based on their median PPIDR values 27.55% vs. 30.85%, whereas median ADS values are almost indistinguishable (0.407 vs. 0.398). In the both cases, the observed difference between the two groups in the median values were not statistically significant (p = 0.193 for median PPIDR and p = 0.788 for median ADS). Figure 9 represents aligned disorder profiles of the S100 proteins and shows that the disorder profiles of the S100 proteins capable of interaction with the cytokines seems to be more similar to each other than the disorder profiles of the S100 proteins that cannot interact with the cytokines (see Figures 9A and 9B, respectively).

Overall, results of these analyses provided some interesting hints on the potential involvement of intrnsic disorder in the S100-cytokine interactions. Our data suggest that the TNF domains of the TNFSF members capable of interaction with S100 proteins are somewhat more disordered than the TNF domains of the non-interacting cytokines, but the TNF domains of the non-interacting cytokines are more similar to each other. On the other hand, the S100 proteins capable of interaction with the cytokines are somewht more ordered than the non-interacting S100 proteins and show more similarity in their disorder profiles, suggesting that some specific disorder-related sequnce peculiarities may play a role in selectivity of the S100-cytokine interactions.

## 4. Conclusions

Among soluble forms of the 13 representatives of TNFSF family studied in this work and 3 TNFSF members previously studied [17,18] for their specificity to non-fused S100 proteins (Table S1), 10 cytokines demonstrated the ability to bind up to 8 S100 proteins, with a total number of positive interactions reaching 31, and the corresponding dissociation constants estimated by SPR spectroscopy ranged from 2 nM to 24 μM (Table 1, Figures 4-5 and ref. [17,18]). The idea of existence of the highly intertwined network of the cytokine-S100 interactions is illustrated by Figure 10 that assembles in one plot all the interaction data known so far. It is obvious that most of the TNFSF members (with the exception to VEGI, CD70, and 4-1BBL) are able to bind more than one S100 protein each. Similarly, most 100 proteins (with the exception to S100A11, S100A13, and S100A16) interact with more than one cytokine.

**Figure 10.**
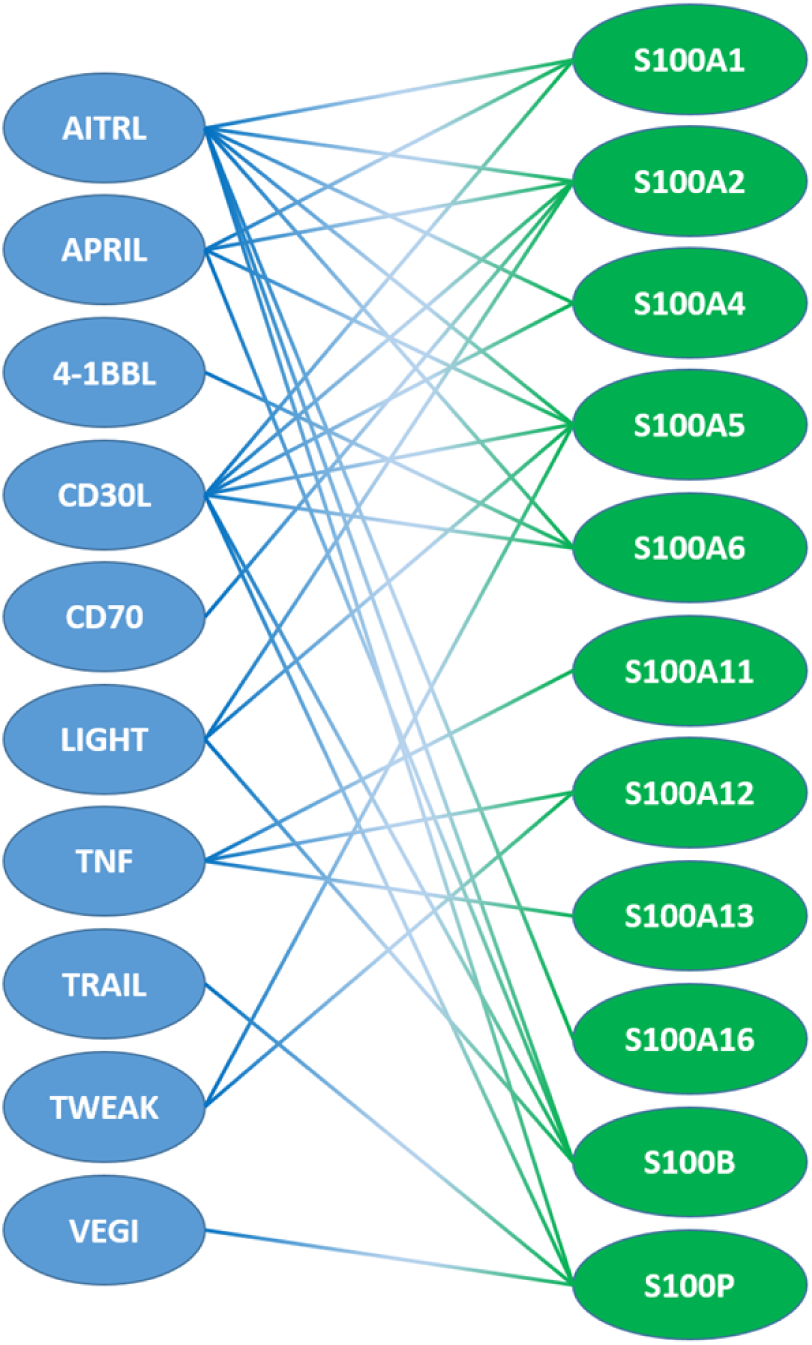
Protein-protein interaction network between soluble forms of the tumor necrosis factor superfamily members and small S100 proteins. This network is assembled based on the data reported in this study and previous studies [17,18].

Overall, about 9% of the TNFSF-S100 interactions examined are positive, and *ca* 63% of the TNFSF members specifically bind at least one S100 protein, which is similar to the 71-73% of the four-helical cytokines shown to be specific to S100A6/S100P [33,57]. Molecular docking and partial experimental verification of the theoretical predictions evidence that S100 proteins bind TNFSF and four-helical cytokines in a similar manner, characteristic for target recognition by S100 proteins. Although we found some weak correlations between the disorder predispositions of the members of the TNFSF superfamily and S100 proteins and their capability to be involved in mutual interactions, structural reasons for the selectivity of the TNFSF-S100 interactions remain unclear, highlighting the need for direct structural studies of this aspect.

The ability of certain S100 proteins to recognize multiple TNFSF members reflects the inherent potential of S100 family to share binding targets, as demonstrated by correlation of the number of TNFSF members bound per an S100 protein with the “promiscuity parameter” [71] (Figure 5B). Meanwhile, some of the S100 proteins with lowest promiscuity parameter (namely, S100A11/A12/A13) exhibit highest affinity to TNFSF members, suggesting a possible (patho)physiological significance of these selective interactions. This assumption is supported by results of cellular studies in the case of S100A12/A13 binding to TNF [18]. Similarly, TRAIL-S100P interaction shows signs of physiological importance in the cellular assays [17] despite much higher promiscuity parameter of S100P and moderate values of the dissociation constant. This fact suggests that other TNFSF-S100 interactions may have physiological significance under certain conditions, particularly in the extracellular milieu of damaged cells.

Since the interactions of S100 proteins with TNFSF representatives characterized to date inhibit activity of the latter [17,18], it is likely that the identified TNFSF-S100 interactions serve as a mechanism for suppression of extracellular signaling in the case of cell damage. Considering that the disease spectra associated with the both protein families overlap significantly (cancer, autoimmune and inflammatory diseases), the TNFSF-S100 interactions are attractive for targeted pharmacological interventions modulating these interactions. Furthermore, S100 protein levels may need to be considered upon use and development of the drugs targeting TNFSF member signaling to ensure their maximum efficacy.

## Abbreviations

4-1BBL: 4-1BB ligand (TNF ligand superfamily member 9)
ADS: average disorder score
AITRL: Activation-inducible TNF-related ligand (TNF ligand superfamily member 18)
APRIL: A proliferation-inducing ligand (TNF ligand superfamily member 13)
BAFF: B-cell-activating factor (TNF ligand superfamily member 13B)
BCMA: B-cell maturation protein
CaM: calmodulin
CD30L: CD30 ligand (TNF ligand superfamily member 8)
CD70: CD70 antigen (TNF ligand superfamily member 7)
DR: Death receptor
EDAC: N-(3-dimethylaminopropyl)-N ′ -ethylcarbodiimide hydrochloride
EDTA: ethylenediaminetetraacetic acid
HEPES: 4-(2-hydroxyethyl)piperazine-1-ethanesulfonic acid
HSA: human serum albumin
IFN: interferon
LIGHT: TNF ligand superfamily member 14
NMR: nuclear magnetic resonance
PDB: Protein Data Bank [62]
PPIDR: percent of intrinsically disordered residues
PV: α-parvalbumin
RAGE: receptor for advanced glycation end products
RANKL: Receptor activator of nuclear factor κ-B
RU: resonance unit
SPR: surface plasmon resonance
sulfo-NHS: N-hydroxysulfosuccinimide sodium salt
THD: TNF homology domain
TNF: Tumor necrosis factor (TNF ligand superfamily member 2)
TNFSF: TNF superfamily
TRAIL: TNF-related apoptosis-inducing ligand (TNF ligand superfamily member 10)
TWEAK: TNF-related weak inducer of apoptosis (TNF ligand superfamily member 12)
VEGI: Vascular endothelial cell growth inhibitor (TNF ligand superfamily member 15)
Δ42–47: human S100P mutant lacking PGFLQS sequence in the ‘hinge’ region [57].

## Supplementary Materials

The following supporting information can be downloaded at: www.mdpi.com/xxx/s1: Supplementary Materials SM1: Data for Figure 6. Figure S1: SPR spectroscopy data on kinetics of interaction between the TNFSF members immobilized on SPR sensor by amine coupling and S100 proteins or human CaM/PV/HSA (1 μM) at 25°C, in the absence/presence of Ca^2+^; Figure S2: The predicted contact residues of the TNFSF-S100 complexes mapped onto the amino acid sequences of the binding partners aligned with Clustal Omega; Table S1: The samples of soluble forms of TNFSF representatives studied in this work and refs. [17,18] for affinity to non-fused S100 proteins and their characteristics; Table S2: The structures of the Ca^2+^-loaded S100 dimers and TNFSF members used for structural modelling of their complexes using ClusPro docking server; Table S3: The contact residues of the complexes between Ca^2+^-loaded S100 dimers and TNFSF members predicted using ClusPro docking server.

## Author Contributions

Conceptualization, S.E.P.; validation, V.A.R., A.S.K., E.I.D., V.N.U., S.E.P.; formal analysis, V.A.R., E.I.D., V.N.U., S.E.P.; investigation, V.A.R., A.S.K., E.I.D., A.S.S., M.E.P., E.A.L., V.N.U.; data curation, V.A.R., E.I.D., V.N.U., S.E.P.; writing—original draft preparation, V.A.R., E.I.D., V.N.U., S.E.P.; writing—review and editing, V.N.U., E.A.P., S.E.P.; supervision, S.E.P.; project administration, S.E.P.; funding acquisition, S.E.P. All authors have read and agreed to the published version of the manuscript.

## Funding

This research was funded by Russian Science Foundation, grant № 25-14-00223 to S.E.P.

## Institutional Review Board Statement

Not applicable.

## Informed Consent Statement

Not applicable.

## Data Availability Statement

The data present in the current study are available from the corresponding authors on reasonable request.

## Conflicts of Interest

The authors declare no conflict of interest. The funders had no role in the design of the study; in the collection, analyses, or interpretation of data; in the writing of the manuscript; or in the decision to publish the results.

## Notes

### Competing Interest Statement

The authors have declared no competing interest.

## References

1. Kolfschoten, G.M.; Pradet-Balade, B.; Hahne, M.; Medema, J.P. TWE-PRIL; a fusion protein of TWEAK and APRIL. Biochem Pharmacol 2003, 66, 1427–1432, doi:10.1016/s0006-2952(03)00493-3.

2. Dostert, C.; Grusdat, M.; Letellier, E.; Brenner, D. The TNF Family of Ligands and Receptors: Communication Modules in the Immune System and Beyond. Physiol Rev 2019, 99, 115–160, doi:10.1152/physrev.00045.2017.

3. Bodmer, J.L.; Schneider, P.; Tschopp, J. The molecular architecture of the TNF superfamily. Trends Biochem Sci 2002, 27, 19–26, doi:10.1016/s0968-0004(01)01995-8.

4. Locksley, R.M.; Killeen, N.; Lenardo, M.J. The TNF and TNF receptor superfamilies: integrating mammalian biology. Cell 2001, 104, 487–501, doi:10.1016/s0092-8674(01)00237-9.

5. Carswell, E.A.; Old, L.J.; Kassel, R.L.; Green, S.; Fiore, N.; Williamson, B. An endotoxin-induced serum factor that causes necrosis of tumors. Proc Natl Acad Sci U S A 1975, 72, 3666–3670, doi:10.1073/pnas.72.9.3666.

6. Lu, J.; Hu, D.; Zhang, Y.; Ma, C.; Shen, L.; Shuai, B. Current comprehensive understanding of denosumab (the RANKL neutralizing antibody) in the treatment of bone metastasis of malignant tumors, including pharmacological mechanism and clinical trials. Front Oncol 2023, 13, 1133828, doi:10.3389/fonc.2023.1133828.

7. Kaegi, C.; Steiner, U.C.; Wuest, B.; Crowley, C.; Boyman, O. Systematic review of safety and efficacy of belimumab in treating immune-mediated disorders. Allergy 2021, 76, 2673–2683, doi:10.1111/all.14704.

8. Zeng, L.; Yang, K.; Wu, Y.; Yu, G.; Yan, Y.; Hao, M.; Song, T.; Li, Y.; Chen, J.; Sun, L. Telitacicept: A novel horizon in targeting autoimmunity and rheumatic diseases. J Autoimmun 2024, 148, 103291, doi:10.1016/j.jaut.2024.103291.

9. Jiang, Y.; Dong, S.; Wang, Y. Antibody-Drug Conjugates Targeting CD30 in T-Cell Lymphomas: Clinical Progression and Mechanism. Cancers (Basel) 2025, 17, doi:10.3390/cancers17030496.

10. Mukhopadhyay, P.; Abdullah, H.A.; Opalinska, J.B.; Paka, P.; Richards, E.; Weisel, K.; Trudel, S.; Mateos, M.V.; Dimopoulos, M.A.; Lonial, S. The clinical journey of belantamab mafodotin in relapsed or refractory multiple myeloma: lessons in drug development. Blood Cancer J 2025, 15, 15, doi:10.1038/s41408-025-01212-0.

11. Croft, M.; Salek-Ardakani, S.; Ware, C.F. Targeting the TNF and TNFR superfamilies in autoimmune disease and cancer. Nat Rev Drug Discov 2024, 23, 939–961, doi:10.1038/s41573-024-01053-9.

12. Veerasubramanian, P.K.; Wynn, T.A.; Quan, J.; Karlsson, F.J. Targeting TNF/TNFR superfamilies in immune-mediated inflammatory diseases. J Exp Med 2024, 221, doi:10.1084/jem.20240806.

13. Ababneh, O.; Nishizaki, D.; Kato, S.; Kurzrock, R. Tumor necrosis factor superfamily signaling: life and death in cancer. Cancer Metastasis Rev 2024, 43, 1137–1163, doi:10.1007/s10555-024-10206-6.

14. Chandonia, J.M.; Guan, L.; Lin, S.; Yu, C.; Fox, N.K.; Brenner, S.E. SCOPe: improvements to the structural classification of proteins - extended database to facilitate variant interpretation and machine learning. Nucleic Acids Res 2022, 50, D553–D559, doi:10.1093/nar/gkab1054.

15. Ross, E.; Ali, M.; Woo, L.; Chadwick, S.; Lee, E.; Sangston, R.; Sood, C.; Zhang, Q.; Zhang, X.; Deppmann, C. Reverse Signalling. 2016; pp. 1–7.

16. Dhusia, K.; Su, Z.; Wu, Y. Computational analyses of the interactome between TNF and TNFR superfamilies. Comput Biol Chem 2023, 103, 107823, doi:10.1016/j.compbiolchem.2023.107823.

17. Rastrygina, V.A.; Kazakov, A.S.; Fadeev, R.S.; Meshcheriakova, E.I.; Deryusheva, E.I.; Sokolov, A.S.; Permyakova, M.E.; Litus, E.A.; Uversky, V.N.; Permyakov, E.A.;, et al. Soluble form of tumor necrosis factor-related apoptosis-inducing ligand interacts with S100P protein. Int J Biol Macromol 2025, 143667, doi:10.1016/j.ijbiomac.2025.143667.

18. Kazakov, A.S.; Zemskova, M.Y.; Rystsov, G.K.; Vologzhannikova, A.A.; Deryusheva, E.I.; Rastrygina, V.A.; Sokolov, A.S.; Permyakova, M.E.; Litus, E.A.; Uversky, V.N.;, et al. Specific S100 Proteins Bind Tumor Necrosis Factor and Inhibit Its Activity. Int J Mol Sci 2022, 23, 15956, doi:10.3390/ijms232415956.

19. Donato, R. S100: a multigenic family of calcium-modulated proteins of the EF-hand type with intracellular and extracellular functional roles. Int J Biochem Cell Biol 2001, 33, 637–668, doi:S1357-2725(01)00046-2 [pii].

20. Donato, R.; Cannon, B.R.; Sorci, G.; Riuzzi, F.; Hsu, K.; Weber, D.J.; Geczy, C.L. Functions of S100 Proteins. Curr Mol Med 2013, 13, 24–57.

21. Sreejit, G.; Flynn, M.C.; Patil, M.; Krishnamurthy, P.; Murphy, A.J.; Nagareddy, P.R. S100 family proteins in inflammation and beyond. Adv Clin Chem 2020, 98, 173–231, doi:10.1016/bs.acc.2020.02.006.

22. Singh, P.; Ali, S.A. Multifunctional Role of S100 Protein Family in the Immune System: An Update. Cells 2022, 11, 2274, doi:10.3390/cells11152274.

23. Gonzalez, L.L.; Garrie, K.; Turner, M.D. Role of S100 proteins in health and disease. Biochim Biophys Acta Mol Cell Res 2020, 1867, 118677, doi:10.1016/j.bbamcr.2020.118677.

24. Fritz, G.; Heizmann, C.W. 3D Structures of the Calcium and Zinc Binding S100 Proteins. In Handbook of Metalloproteins, John Wiley & Sons, L., Ed.; John Wiley & Sons: Hoboken, NJ, USA, 2004.

25. Nockolds, C.E.; Kretsinger, R.H.; Coffee, C.J.; Bradshaw, R.A. Structure of a calcium-binding carp myogen. Proc Natl Acad Sci U S A 1972, 69, 581–584.

26. Gilston, B.A.; Skaar, E.P.; Chazin, W.J. Binding of transition metals to S100 proteins. Sci China Life Sci 2016, 59, 792–801, doi:10.1007/s11427-016-5088-4.

27. Moroz, O.V.; Wilson, K.S.; Bronstein, I.B. The role of zinc in the S100 proteins: insights from the X-ray structures. Amino Acids 2010, doi:10.1007/s00726-010-0540-4.

28. UniProt, C. UniProt: the Universal Protein Knowledgebase in 2023. Nucleic Acids Res 2023, 51, D523–D531, doi:10.1093/nar/gkac1052.

29. Signor, L.; Paris, T.; Mas, C.; Picard, A.; Lutfalla, G.; Boeri Erba, E.; Yatime, L. Divalent cations influence the dimerization mode of murine S100A9 protein by modulating its disulfide bond pattern. J Struct Biol 2021, 213, 107689, doi:10.1016/j.jsb.2020.107689.

30. Winningham-Major, F.; Staecker, J.L.; Barger, S.W.; Coats, S.; Van Eldik, L.J. Neurite extension and neuronal survival activities of recombinant S100 beta proteins that differ in the content and position of cysteine residues. J Cell Biol 1989, 109, 3063–3071.

31. Fritz, G.; Botelho, H.M.; Morozova-Roche, L.A.; Gomes, C.M. Natural and amyloid self-assembly of S100 proteins: structural basis of functional diversity. FEBS J 2010, 277, 4578–4590, doi:10.1111/j.1742-4658.2010.07887.x.

32. Santamaria-Kisiel, L.; Rintala-Dempsey, A.C.; Shaw, G.S. Calcium-dependent and -independent interactions of the S100 protein family. Biochem J 2006, 396, 201–214, doi:10.1042/BJ20060195.

33. Kazakov, A.S.; Deryusheva, E.I.; Rastrygina, V.A.; Sokolov, A.S.; Permyakova, M.E.; Litus, E.A.; Uversky, V.N.; Permyakov, E.A.; Permyakov, S.E. Interaction of S100A6 Protein with the Four-Helical Cytokines. Biomolecules 2023, 13, 1345.

34. Hermann, A.; Donato, R.; Weiger, T.M.; Chazin, W.J. S100 calcium binding proteins and ion channels. Front Pharmacol 2012, 3, doi:10.3389/Fphar.2012.00067.

35. Marenholz, I.; Volz, A.; Ziegler, A.; Davies, A.; Ragoussis, I.; Korge, B.P.; Mischke, D. Genetic analysis of the epidermal differentiation complex (EDC) on human chromosome 1q21: chromosomal orientation, new markers, and a 6-Mb YAC contig. Genomics 1996, 37, 295–302, doi:10.1006/geno.1996.0563.

36. Solomon, E.; Borrow, J.; Goddard, A.D. Chromosome aberrations and cancer. Science 1991, 254, 1153–1160, doi:10.1126/science.1957167.

37. Abdi, W.; Romasco, A.; Alkurdi, D.; Santacruz, E.; Okinedo, I.; Zhang, Y.; Kannan, S.; Shakiba, S.; Richmond, J.M. An overview of S100 proteins and their functions in skin homeostasis, interface dermatitis conditions and other skin pathologies. Exp Dermatol 2024, 33, e15158, doi:10.1111/exd.15158.

38. Bresnick, A.R.; Weber, D.J.; Zimmer, D.B. S100 proteins in cancer. Nat Rev Cancer 2015, 15, 96–109, doi:10.1038/nrc3893.

39. Allgower, C.; Kretz, A.L.; von Karstedt, S.; Wittau, M.; Henne-Bruns, D.; Lemke, J. Friend or Foe: S100 Proteins in Cancer. Cancers 2020, 12, 2037, doi:10.3390/cancers12082037.

40. Liang, X.; Huang, X.; Cai, Z.; Deng, Y.; Liu, D.; Hu, J.; Jin, Z.; Zhou, X.; Zhou, H.; Wang, L. The S100 family is a prognostic biomarker and correlated with immune cell infiltration in pan-cancer. Discov Oncol 2024, 15, 137, doi:10.1007/s12672-024-00945-x.

41. Hua, X.; Zhang, H.M.; Jia, J.F.; Chen, S.S.; Sun, Y.; Zhu, X.L. Roles of S100 family members in drug resistance in tumors: Status and prospects. Biomed Pharmacother 2020, 127, doi:10.1016/J.Biopha.2020.110156.

42. Manfredi, M.; Van Hoovels, L.; Benucci, M.; De Luca, R.; Coccia, C.; Bernardini, P.; Russo, E.; Amedei, A.; Guiducci, S.; Grossi, V.;, et al. Circulating Calprotectin (cCLP) in autoimmune diseases. Autoimmun Rev 2023, 22, 103295, doi:10.1016/j.autrev.2023.103295.

43. Li, W.; Chen, Q.; Peng, C.; Yang, D.; Liu, S.; Lv, Y.; Jiang, L.; Xu, S.; Huang, L. Roles of the Receptor for Advanced Glycation End Products and Its Ligands in the Pathogenesis of Alzheimer’s Disease. Int J Mol Sci 2025, 26, doi:10.3390/ijms26010403.

44. Sattar, Z.; Lora, A.; Jundi, B.; Railwah, C.; Geraghty, P. The S100 Protein Family as Players and Therapeutic Targets in Pulmonary Diseases. Pulm Med 2021, 2021, 5488591, doi:10.1155/2021/5488591.

45. Cheng, B.; Bian, Y.; Song, X.; Li, W.; Li, M.; Feng, R. Role of S100A1, S100A4, S100A8/A9 and S100B in myocardial infarction and heart failure. Int Immunopharmacol 2025, 151, 114348, doi:10.1016/j.intimp.2025.114348.

46. Zhou, Y.; Zha, Y.; Yang, Y.; Ma, T.; Li, H.; Liang, J. S100 proteins in cardiovascular diseases. Mol Med 2023, 29, 68, doi:10.1186/s10020-023-00662-1.

47. Xia, P.; Ji, X.; Yan, L.; Lian, S.; Chen, Z.; Luo, Y. Roles of S100A8, S100A9 and S100A12 in infection, inflammation and immunity. Immunology 2024, 171, 365-376, doi:10.1111/imm.13722.

48. Bresnick, A.R. S100 proteins as therapeutic targets. Biophys Rev 2018, 10, 1617–1629, doi:10.1007/s12551-018-0471-y.

49. Kurpet, K.; Chwatko, G. S100 Proteins as Novel Therapeutic Targets in Psoriasis and Other Autoimmune Diseases. Molecules 2022, 27, doi:10.3390/molecules27196640.

50. Frauchiger, A.L.; Dummer, R.; Mangana, J. Serum S100B Levels in Melanoma. Methods Mol Biol 2019, 1929, 691–700, doi:10.1007/978-1-4939-9030-6_43.

51. Sejersen, K.; Eriksson, M.B.; Larsson, A.O. Calprotectin as a Biomarker for Infectious Diseases: A Comparative Review with Conventional Inflammatory Markers. Int J Mol Sci 2025, 26, doi:10.3390/ijms26136476.

52. Kazakov, A.S.; Sokolov, A.S.; Vologzhannikova, A.A.; Permyakova, M.E.; Khorn, P.A.; Ismailov, R.G.; Denessiouk, K.A.; Denesyuk, A.I.; Rastrygina, V.A.; Baksheeva, V.E.;, et al. Interleukin-11 binds specific EF-hand proteins via their conserved structural motifs. J Biomol Struct Dyn 2017, 35, 78–91, doi:10.1080/07391102.2015.1132392.

53. Kazakov, A.S.; Sofin, A.D.; Avkhacheva, N.V.; Denesyuk, A.I.; Deryusheva, E.I.; Rastrygina, V.A.; Sokolov, A.S.; Permyakova, M.E.; Litus, E.A.; Uversky, V.N.;, et al. Interferon Beta Activity Is Modulated via Binding of Specific S100 Proteins. International journal of molecular sciences 2020, 21, 9473, doi:10.3390/ijms21249473.

54. Kazakov, A.S.; Deryusheva, E.I.; Sokolov, A.S.; Permyakova, M.E.; Litus, E.A.; Rastrygina, V.A.; Uversky, V.N.; Permyakov, E.A.; Permyakov, S.E. Erythropoietin Interacts with Specific S100 Proteins. Biomolecules 2022, 12, 120, doi:10.3390/biom12010120.

55. Kazakov, A.S.; Rastrygina, V.A.; Vologzhannikova, A.A.; Zemskova, M.Y.; Bobrova, L.A.; Deryusheva, E.I.; Permyakova, M.E.; Sokolov, A.S.; Litus, E.A.; Shevelyova, M.P.;, et al. Recognition of granulocyte-macrophage colony-stimulating factor by specific S100 proteins. Cell Calcium 2024, 119, 102869, doi:10.1016/j.ceca.2024.102869.

56. Kazakov, A.S.; Sokolov, A.S.; Rastrygina, V.A.; Solovyev, V.V.; Ismailov, R.G.; Mikhailov, R.V.; Ulitin, A.B.; Yakovenko, A.R.; Mirzabekov, T.A.; Permyakov, E.A.;, et al. High-affinity interaction between interleukin-11 and S100P protein. Biochem. Biophys. Res. Commun. 2015, 468, 733–738, doi:10.1016/j.bbrc.2015.11.024.

57. Kazakov, A.S.; Deryusheva, E.I.; Permyakova, M.E.; Sokolov, A.S.; Rastrygina, V.A.; Uversky, V.N.; Permyakov, E.A.; Permyakov, S.E. Calcium-Bound S100P Protein Is a Promiscuous Binding Partner of the Four-Helical Cytokines. Int J Mol Sci 2022, 23, 12000, doi:10.3390/ijms231912000.

58. Gopalakrishna, R.; Anderson, W.B. Ca2+-induced hydrophobic site on calmodulin: application for purification of calmodulin by phenyl-Sepharose affinity chromatography. Biochem. Biophys. Res. Commun. 1982, 104, 830–836, doi:10.1016/0006-291x(82)90712-4.

59. Vologzhannikova, A.A.; Shevelyova, M.P.; Kazakov, A.S.; Sokolov, A.S.; Borisova, N.I.; Permyakov, E.A.; Kircheva, N.; Nikolova, V.; Dudev, T.; Permyakov, S.E. Strontium Binding to alpha-Parvalbumin, a Canonical Calcium-Binding Protein of the "EF-Hand" Family. Biomolecules 2021, 11, doi:10.3390/biom11081158.

60. Pace, C.N.; Vajdos, F.; Fee, L.; Grimsley, G.; Gray, T. How to measure and predict the molar absorption coefficient of a protein. Protein Sci. 1995, 4, 2411–2423.

61. Desta, I.T.; Porter, K.A.; Xia, B.; Kozakov, D.; Vajda, S. Performance and Its Limits in Rigid Body Protein-Protein Docking. Structure 2020, 28, 1071–1081 e1073, doi:10.1016/j.str.2020.06.006.

62. Berman, H.M.; Westbrook, J.; Feng, Z.; Gilliland, G.; Bhat, T.N.; Weissig, H.; Shindyalov, I.N.; Bourne, P.E. The Protein Data Bank. Nucleic Acids Res. 2000, 28, 235–242.

63. Abramson, J.; Adler, J.; Dunger, J.; Evans, R.; Green, T.; Pritzel, A.; Ronneberger, O.; Willmore, L.; Ballard, A.J.; Bambrick, J.;, et al. Accurate structure prediction of biomolecular interactions with AlphaFold 3. Nature 2024, 630, 493–500, doi:10.1038/s41586-024-07487-w.

64. Madeira, F.; Madhusoodanan, N.; Lee, J.; Eusebi, A.; Niewielska, A.; Tivey, A.R.N.; Lopez, R.; Butcher, S. The EMBL-EBI Job Dispatcher sequence analysis tools framework in 2024. Nucleic Acids Res 2024, 52, W521–W525, doi:10.1093/nar/gkae241.

65. Schrodinger, LLC. The PyMOL Molecular Graphics System, Version 1.8. 2015.

66. Lang, I.; Fullsack, S.; Wyzgol, A.; Fick, A.; Trebing, J.; Arana, J.A.; Schafer, V.; Weisenberger, D.; Wajant, H. Binding Studies of TNF Receptor Superfamily (TNFRSF) Receptors on Intact Cells. J Biol Chem 2016, 291, 5022–5037, doi:10.1074/jbc.M115.683946.

67. Leclerc, E.; Fritz, G.; Vetter, S.W.; Heizmann, C.W. Binding of S100 proteins to RAGE: an update. Biochim Biophys Acta 2009, 1793, 993–1007, doi:10.1016/j.bbamcr.2008.11.016.

68. Yap, K.L.; Kim, J.; Truong, K.; Sherman, M.; Yuan, T.; Ikura, M. Calmodulin target database. J Struct Funct Genomics 2000, 1, 8–14, doi:10.1023/a:1011320027914.

69. Ferdinand, M. Polyamino carboxylic acids and process of making same. US2130505A, 1938.

70. Kazakov, A.S.; Sokolov, A.S.; Permyakova, M.E.; Litus, E.A.; Uversky, V.N.; Permyakov, E.A.; Permyakov, S.E. Specific cytokines of interleukin-6 family interact with S100 proteins. Cell Calcium 2022, 101, 102520, doi:10.1016/j.ceca.2021.102520.

71. Simon, M.A.; Bartus, É.; Mag, B.; Boros, E.; Roszjár, L.; Gógl, G.; Travé, G.; Martinek, T.A.; Nyitray, L. Promiscuity mapping of the S100 protein family using a high-throughput holdup assay. Sci Rep 2022, 12, 5904, doi:10.1038/s41598-022-09574-2.

72. Simon, M.A.; Ecsedi, P.; Kovacs, G.M.; Poti, A.L.; Remenyi, A.; Kardos, J.; Gogl, G.; Nyitray, L. High-throughput competitive fluorescence polarization assay reveals functional redundancy in the S100 protein family. The FEBS journal 2020, 287, 2834–2846, doi:10.1111/febs.15175.

73. Del Toro, N.; Shrivastava, A.; Ragueneau, E.; Meldal, B.; Combe, C.; Barrera, E.; Perfetto, L.; How, K.; Ratan, P.; Shirodkar, G.;, et al. The IntAct database: efficient access to fine-grained molecular interaction data. Nucleic Acids Res 2022, 50, D648–D653, doi:10.1093/nar/gkab1006.

74. Oughtred, R.; Rust, J.; Chang, C.; Breitkreutz, B.J.; Stark, C.; Willems, A.; Boucher, L.; Leung, G.; Kolas, N.; Zhang, F.;, et al. The BioGRID database: A comprehensive biomedical resource of curated protein, genetic, and chemical interactions. Protein Sci 2021, 30, 187–200, doi:10.1002/pro.3978.

75. Deutsch, E.W. The PeptideAtlas Project. Methods Mol Biol 2010, 604, 285–296, doi:10.1007/978-1-60761-444-9_19.

76. Permyakov, S.E.; Ismailov, R.G.; Xue, B.; Denesyuk, A.I.; Uversky, V.N.; Permyakov, E.A. Intrinsic disorder in S100 proteins. Mol Biosyst 2011, 7, 2164–2180, doi:10.1039/c0mb00305k.

77. Penumutchu, S.R.; Chou, R.H.; Yu, C. Structural Insights into Calcium-Bound S100P and the V Domain of the RAGE Complex. Plos One 2014, 9, e103947, doi:10.1371/journal.pone.0103947.

78. Xue, B.; Dunbrack, R.L.; Williams, R.W.; Dunker, A.K.; Uversky, V.N. PONDR-FIT: a meta-predictor of intrinsically disordered amino acids. Biochim Biophys Acta 2010, 1804, 996–1010, doi:10.1016/j.bbapap.2010.01.011.

79. Sievers, F.; Wilm, A.; Dineen, D.; Gibson, T.J.; Karplus, K.; Li, W.; Lopez, R.; McWilliam, H.; Remmert, M.; Soding, J.;, et al. Fast, scalable generation of high-quality protein multiple sequence alignments using Clustal Omega. Mol Syst Biol 2011, 7, 539, doi:10.1038/msb.2011.75.

80. Jumper, J.; Evans, R.; Pritzel, A.; Green, T.; Figurnov, M.; Ronneberger, O.; Tunyasuvunakool, K.; Bates, R.; Zidek, A.; Potapenko, A.;, et al. Highly accurate protein structure prediction with AlphaFold. Nature 2021, 596, 583–589, doi:10.1038/s41586-021-03819-2.

81. Braberg, H.; Webb, B.M.; Tjioe, E.; Pieper, U.; Sali, A.; Madhusudhan, M.S. SALIGN: a web server for alignment of multiple protein sequences and structures. Bioinformatics 2012, 28, 2072–2073, doi:10.1093/bioinformatics/bts302.

